# NetBCE: An Interpretable Deep Neural Network for Accurate Prediction of Linear B-Cell Epitopes

**DOI:** 10.1101/2022.05.23.493092

**Authors:** Haodong Xu, Zhongming Zhao

**Author notes:** Corresponding author. (Zhao Z).

## Abstract

Identification of B-cell epitopes (BCEs) plays an essential role in the development of peptide vaccines, immuno-diagnostic reagents, and antibody design and production. In this work, we generated a large benchmark dataset comprising 126,779 experimentally-supported, linear epitope-containing regions in 3567 protein clusters from over 1.3 million B cell assays. Analysis of this curated dataset showed large pathogen diversity covering 176 different families. The accuracy in linear BCE prediction was found to strongly vary with different features, while the performance by sequence features was superior to that by structural features. To search more efficient and interpretive feature representations, a ten-layer deep learning framework for linear BCE prediction, namely NetBCE, was developed. NetBCE achieved high accuracy and robust performance with the average area under the curve (AUC) value of 0.846 in five-fold cross validation through automatically learning the informative classification features. NetBCE substantially outperformed the conventional machine learning algorithms and other tools, with an over 22.06% improvement of AUC value compared to other tools using an independent dataset. Through investigating the output of important network modules in NetBCE, epitopes and non-epitopes tended to present in distinct regions with efficient feature representation along the network layer hierarchy. The NetBCE tool will be useful for linear B-cell epitopes identification and more generally, immunological and computational biology research.

## Introduction

B-cell epitopes (BCE) represent the regions on antigen surfaces where designated antibodies recognize, bind to, and subsequently induce the immune response in humoral immunity [1, 2]. Identification of B-cell epitopes is a crucial step in immunological studies and medical applications, including peptide-based vaccine development, antibody production and disease prevention [3]. B-cell epitopes are commonly classified into two types: linear epitopes and conformational epitopes. Linear epitopes are composed of a linear sequence of residues from an antigenic sequence, while conformational epitopes refer to atoms on surface residues that come together via protein folding [4]. Many experimental approaches have been developed for BCE identification, including peptide microarrays, X-ray crystallography, and enzyme-linked immunosorbent assay (ELISA) [5]. However, these approaches are expensive and resource intensive. On the other hand, computational approaches have demonstrated promise for predicting linear B-cell epitopes. So far, several computational approaches have been published for linear BCE prediction from protein’s primary sequences [6], including BepiPred [7], iBCE-EL [8], and LBtope [9].

Those initially developed methods typically used and characterized a subset of amino acid physicochemical properties, such as hydrophobicity [10], surface accessibility, flexibility [11] and antigenicity [12]. Later, other feature encoding strategies based on sequence-derived and structural information were applied. These included amino acid composition (AAC), BLOSUM62 scoring matrix, accessible surface area (ASA), secondary structure (SS), and backbone torsion angles (BTA) [13, 14]. However, which features are the most informative for BCE prediction remains unclear. Most of these methods have been developed by using conventional machine-learning algorithms, which may be less powerful in feature representation than deep-learning algorithms [15–18]. Recently, several hundred thousand high-quality linear B-cell epitope assay datasets have been stored in the Immune Epitope Database (IEDB) [19]. This large collection provides a unique opportunity to further develop computational approaches for identification of potential linear B-cell epitope from protein sequences.

In this work, we first collected and curated over 1.3 million B cell assays from the IEDB database. Through quality control procedures, we compiled an experimentally well-characterized dataset, containing more than 126,000 experimentally linear epitope-containing regions from 3567 protein clusters. The curated dataset covered 176 different families, indicating strong pathogen diversity. After homology clearance, we carefully evaluated five types of sequence-derived features [20], six clusters of physicochemical properties [21, 22], as well as three types of structural features [23] using six conventional machine learning (ML) algorithms on the curated dataset. The results show that different types of features displayed various accuracies for linear B cell epitope prediction and sequence features had superior performance compared to structural features. With a sufficient training dataset of B cell assays, the deep neural network can automatically learn informative classification features, making it very appropriate for linear B cell epitope prediction [24]. Therefore, in this study, we developed NetBCE, a ten-layer deep learning framework, and implemented it into tool. The epitope sequences were encoded and taken as input for subsequent feature extraction and representation in the convolution-pooling module. A Bidirectional Long Short-Term Memory (BLSTM) layer was added to retaining features over a long duration and to facilitate the model catching the combinations or dependencies among residues at different positions. Lastly, an attention layer was joined to link the BLSTM layer and the output layer. NetBCE outperformed conventional ML methods by an improvement of the area under curve (AUC) value in a range of 8.77-21.58% using the same training dataset. Moreover, in our comparison of NetBCE with other existing tools using an independent dataset, NetBCE achieved performance with the AUC values of 0.84, and had AUC value improvement by ≥ 22.06% for the linear B cell epitope prediction when compared to other tools. To elucidate the capability of hierarchical representation by NetBCE, we visualized the epitopes and non-epitopes using Uniform Manifold Approximation and Projection (UMAP) [25] based on the feature representation at various network layers. We found that feature representation became more discriminative further along the network layer hierarchy. More specifically, the feature representations for epitopes and non-epitopes sites were mixed at the input layer. As the model continued to train, epitopes and non-epitopes tended to present in distinct regions with efficient feature representation. The NetBCE tool allows the user to explore the data and prediction results in an easily readable and interpretable manner.

## Method

### Data collection and processing

To establish a reliable model, an experimentally-supported dataset was compiled as follows (**Figure 1**). First, we downloaded over 1.3 million B cell assays from the Immune Epitope Database (available at https://www.iedb.org/), the most comprehensive database holding the largest number of experimentally identified epitopes and non-epitopes. Each entry contained an antigen protein sequence with a marked region (hereafter we called “epitope-containing region”) that was an experimentally verified epitope or non-epitope. Protein sequences were retrieved from the National Center for Biotechnology Information (NCBI) [26] and the UniProt database [27] based on the antigen protein IDs provided in the epitope entry. We preprocessed and filtered the dataset by several criteria (Figure 1). First, identical protein sequences were integrated and all related information about epitope-containing regions was aggregated. Second, sequence redundancy for those proteins of non-identical but highly similar was cleared. Using CD-HIT program [28], all proteins were clustered with a threshold of 90% sequence similarity. For each cluster, only the protein having the largest number of epitope-containing regions was retained. To ensure high confidence of the dataset, each epitope assay was carefully explored and regarded as positive hit only when it has been tested as positive in two or more different B-cell assays, whereas those regions that were tested in at least two assays but all were not positive were considered as non-epitopes. In addition, we excluded those candidate epitopes that had less than 5 or more than 25 amino acid residues from the dataset. The number of such epitopes accounted for only a small portion (approximately 1%), but an inclusion of them may result in outliers during model development. Overall, the final non-redundant dataset for training and testing contained 28,714 positive and 98,065 negative epitope-containing regions from 3567 protein sequence clusters, respectively. The compiled dataset was divided into the training dataset (90% of the total epitope-containing regions) and the independent dataset (10% of the remaining epitope-containing regions).

**Figure 1.**
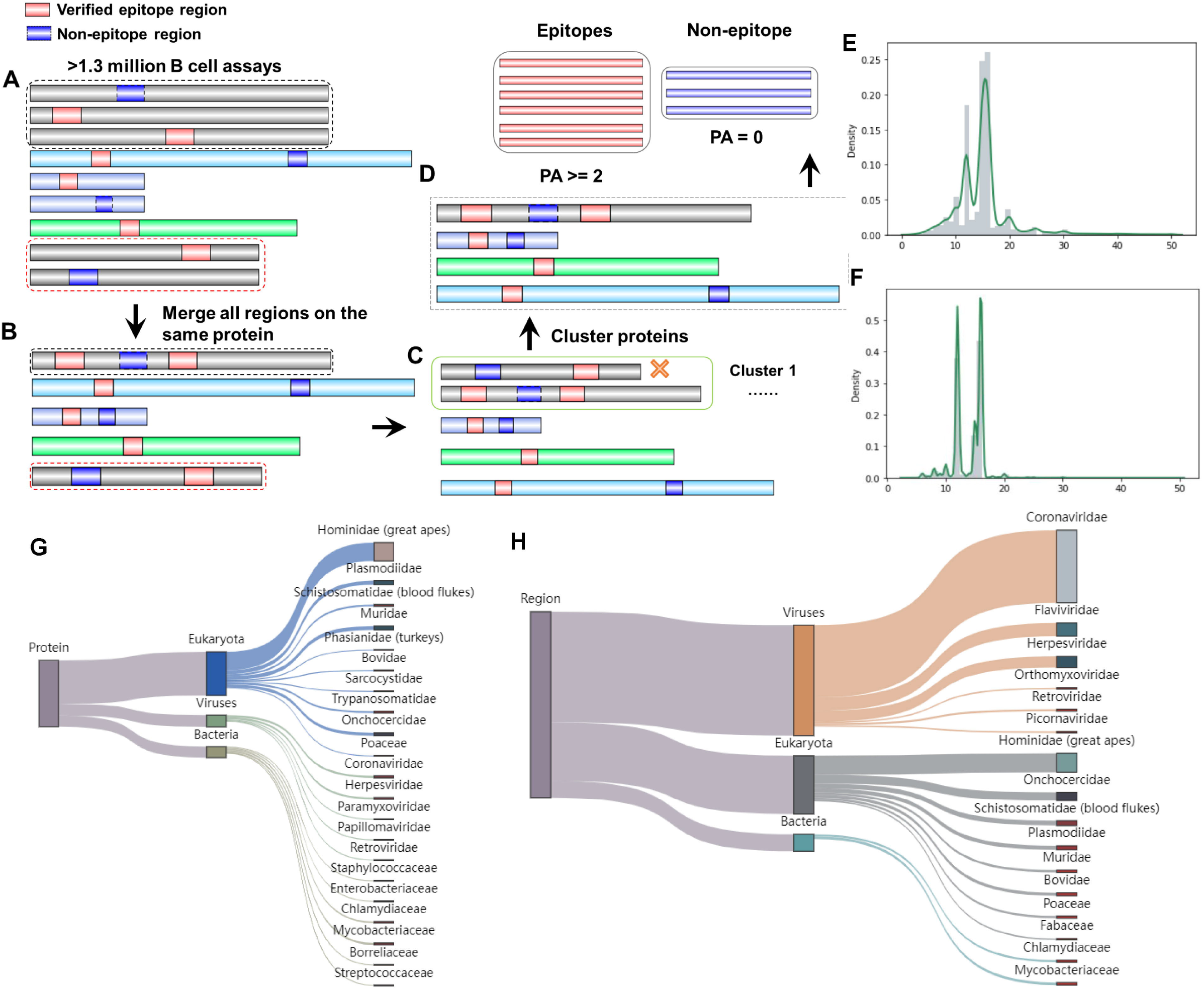
Benchmark data preparation and evaluation. **A.** The experimentally identified epitope-containing regions were collected from the IEDB databases. **B.** Identical protein sequences were integrated and the epitope verified regions were aggregated. **C.** Sequence redundancy was cleaned for the similar proteins by cd-hit. **D.** Proteins with the largest number of epitope-containing regions were retained. The curated dataset was divided into epitopes and non-epitopes according to epitope assay information. The length distribution of (**E**) epitopes and (**F**) non-epitopes. Taxonomic distribution in super-kingdoms and families at the (**G**) protein level and (**H**) verified epitope level.

### Feature encoding

For the curated benchmark dataset, 14 types of features were encoded from the epitope-containing regions of both the positive and negative datasets. These datasets included five types of sequence-derived features, six clusters of physicochemical properties, and three structural features. We classified these 14 feature types as follows. 1) Amino acid composition (AAC). AAC counts the frequencies of 20 types of typical amino acids in epitope-containing regions. 2) Binary. Binary denotes position-specific composition of the amino acids. The 20 types of amino acids were alphabetically sorted and each amino acid was transformed into a binary vector. 3) Composition of K-spaced amino acid pairs (CKSAAP). CKSAAP calculates the composition of amino acid pairs that are separated by *k* other residues within epitope-containing regions. 4) Physicochemical properties, which represent amino acid indices of various physicochemical properties. Numerous studies have indicated strong correlations between physicochemical properties of amino acids and B cell epitopes. In this study, we employed and encoded six categories of properties. They are α and turn propensities, β propensity, hydrophobicity, physicochemical properties, and other properties to characterize these properties. 5) Enhanced amino acid composition (EAAC). EAAC represents the local AAC for the fixed-length sequence window that continuously slides from the 5’ to 3’ terminus of each protein sequence. 6) BLOSUM62 scoring matrix, which is commonly used to score the alignments between evolutionarily divergent protein sequences. 7) ASA. ASA indicates the exposed area of an amino acid residue to solvent. The SPIDER2 tool [23] computes a potential ASA value for each amino acid in epitope-containing regions. 8) SS. SS represents 3 types of structural elements, including α-helix, β-strand, and coil. 9) BTA. BTA measures continuous angle information of the local conformation of proteins, including the backbone torsion angles *φ* and *Ψ*, the angle between Cα_i-1_-Cα_i_-Cα_i+1_ (*θ*) and the dihedral angle rotated about the Cα_i_-Cα_i+1_ bond (*τ*).

### NetBCE model construction

As shown in **Figure 2**, a ten-layer deep learning framework, named NetBCE, was implemented to predict B cell epitopes using amino acid sequences as input. Each layer contained a number of computational units called neurons, which constitutes an internal feature representation. We applied one-hot encoding to convert the epitope sequences to a *m* × 20 binary matrix, where *m* represents the length of the epitope sequence. Then, the binary matrix was entered to a convolution layer [29] to catch sequence sub-motifs. Convolutional kernels act as the crucial components of the convolution layer, which was widely used for sequence motif recognition, regardless of their position in the sequence. A number of studies have used kernels in the convolutional layer to catch sequence patterns from massive sequence data. In the NetBCE, representative patterns were first detected by numerous convolution kernels from the input epitope sequences. The convolutional layer was followed by a maxpooling layer to calculate the maximum activation spots over spatially adjacent regions, and then to summarize the most activated pattern in the sequences. Down sampling strategy in maxpooling downsizes the feature dimension and thus strengthens the deep learning model robustness. To further extract the extensive dependencies of long-range sequence among detected patterns from both forward and backward directions, we added a BLSTM layer [30] in NetBCE. The rationale for adding a BLSTM is that the binding between B cell epitope and B cell receptor may be regulated by multiple spaced amino acids. The power of LSTM for retaining features from a long duration facilitates the model to capture the combinations or dependencies among residues at different positions. The unit in LSTM contains four parts: three gates (input, forget and output) and a single cell remembering features over arbitrary intervals. Specifically, considering an epitope sequence with length L as input 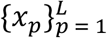 in LSTM, and for every position *p*, denote the input gate as *I_p_*, forget gate as *F_p_*, output gate as *O_p_*, hidden state as *H_p_*, and cell state as *C_p_*. The process of LSTM training is as follows:

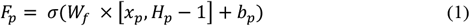

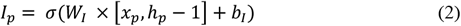

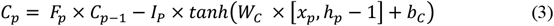

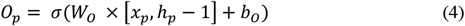

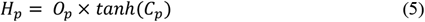

**Figure 2.**
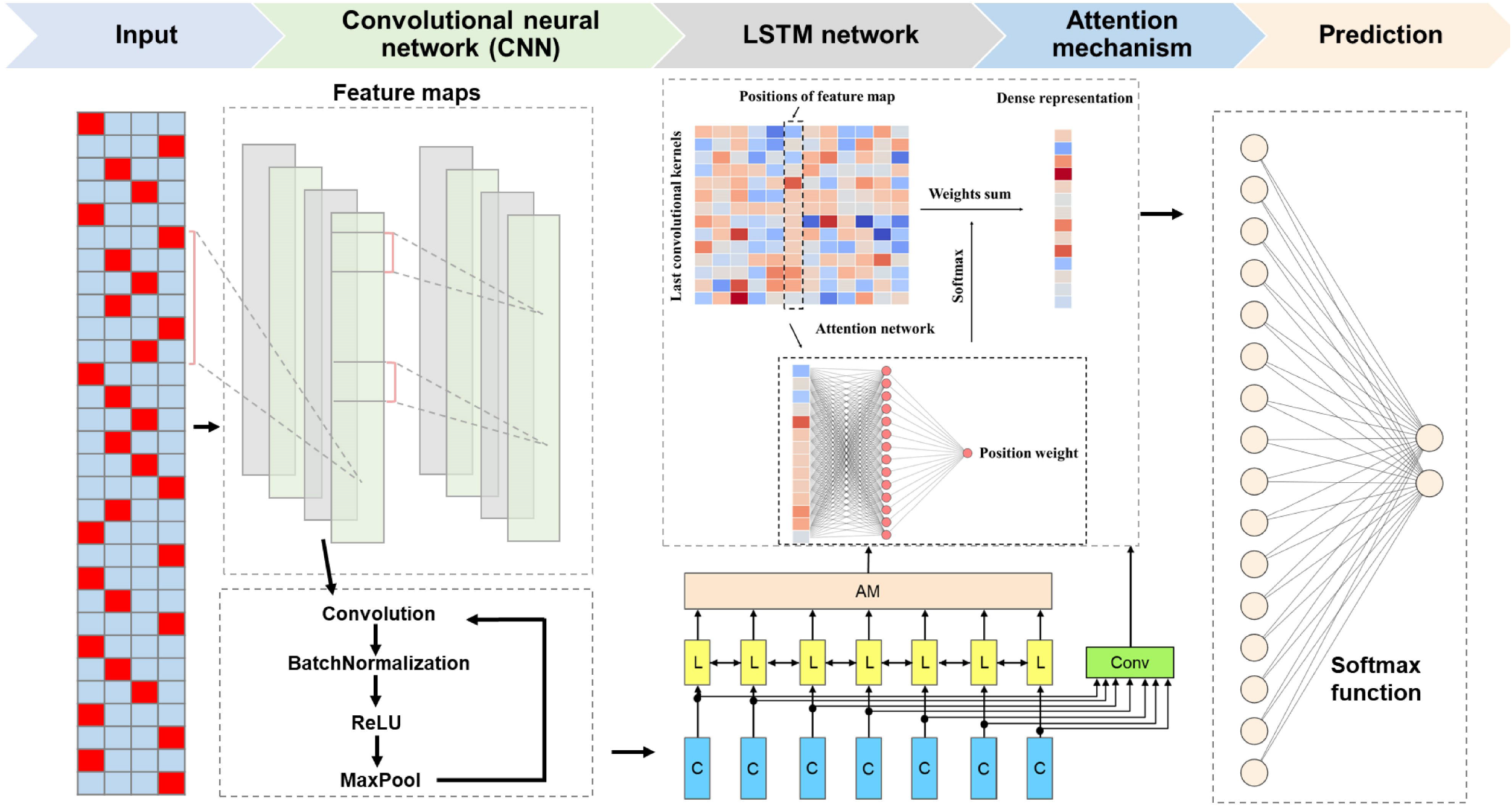
Deep learning framework of NetBCE. NetBCE is built on a ten-layer deep learning framework. The epitope sequences were encoded as binary matrix and taken as input. Then, convolution-pooling module was used for feature extraction and representation. Bidirectional Long Short-Term Memory (BLSTM) layer was added for retaining features from a long duration to capture the combinations or dependencies among residues at different positions. A fully connected layer was used to integrate the variables output from the attention layer and learn the nonlinear relationship. The output layer was composed of two sigmoid neurons for calculating a prediction score for a given peptide.

To further recognize the most representative sequence patterns in NetBCE, an attention layer [31] was added following the BLSTM layer. Because the most distinct patterns may be located somewhere of the epitope, the attention layer was thus adopted to find more informative features by learning the whole hidden states of the BLSTM layer and distribute higher weights to the important locus. Mathematically, by obtaining the hidden variables 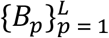 from BLSTM layer as inputs, the attention layer returns the output vector *A* as shown below:

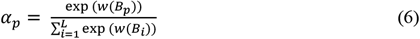

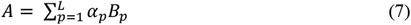

where *w* represents a fully connected neural network that computes a scalar weight.

Finally, we utilized a fully connected layer to integrate the variables output from the attention layer and learn the nonlinear relationships. The output layer was composed of two *sigmoid* neurons calculating a *SBCE* score for a given peptide *y*, as defined as:

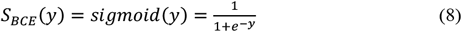

The *SBCE* value, ranging from 0 to 1, represents the probability of peptide to be a real B cell epitope.

### Model training and evaluation

We trained the NetBCE using the Adam optimizer with mini-batch algorithm. The deep learning model was trained to minimize the loss of binary cross-entropy, which catches the difference between the target and predicted label. After each epoch of training, the model was evaluated on the validation dataset, and the corresponding loss and accuracy values were recorded. We introduced an early stop mechanism during training to avoid model overfitting. Specificity, the model was constantly learned until the validation accuracy stopped to increase for twenty epochs. After model training was completed, we evaluated the performance using a test dataset and several metrics were calculated, including accuracy (*Acc*), sensitivity (*Sn*), specificity (*Sp*) and the area under the receiver operating characteristic (ROC) curve (AUC).

The hyperparameters of NetBCE model were optimized to achieve optimal performance using Hyperopt tool [32] via Bayesian mechanism from a list of multiple parameters, including the number of convolutional filters, kernel size, the learning rate, degree of momentum, mini-batch size, strength of parameter regularization, and dropout probability. Hyperopt optimizes the hyperparameter space by creating a classification model upon the metric of the objective function. The probability model was updated after each evaluation of the objective function by incorporating new results. Specifically, 100 evaluations were executed using separate training (inner loop) and validation sets (outer loop). The performance of each set of parameters was evaluated and the corresponding AUC values were calculated. We selected the group of parameters with the highest AUC values as the final parameters of the model. NVIDIA Tensor Cores with four Tesla V100 were used. The Keras version 2.3, a highly useful neural networks API, and the tensorflow-gpu 1.15 version were adopted for a rapid parallel computing.

### Conventional ML classifiers

In this study, we implemented 84 classical machine learning models for prediction of B cell epitope based on nine types of features using six algorithms: AdaBoost (AB), Decision Trees (DT), Gradient Boosting (GB), K-Nearest Neighbors (KNN), Logistic Regression (LR), and Random Forests (RF). Five-fold cross-validation (CV) was performed for each classifier to evaluate the predictive capacity. The ROC curves were illustrated for *Sn vs. 1-Sp* scores and the AUC values were subsequently calculated. For accurate estimation of the performance, the five-fold CV was independently performed by 10 times and the average AUC values was calculated for each model setting. To determine the best parameters for each model, we tested dozens or hundreds of different parameter combinations for each model, and selected the optimal parameters through multiple CVs evaluations.

## Results

### The curated dataset contains large pathogen diversity

From the IEDB database, we extracted over 1.3 million B cell assays with experimentally verified epitope-containing information (**Figure 1**A). After merging all the identical protein sequences, we obtained 8437 proteins preserving 213,700 verified epitope-containing regions (Figure 1B). After removing the redundancy by cd-hit software, 3567 protein sequence clusters were identified. This procedure reduced the number of epitope-containing regions by 40.67% to 126,779 (Figure 1C). By applying our quality control procedures, the final filtered dataset contained 3567 proteins with 28,714 epitopes and 98,065 non-epitopes for model construction (Figure 1D). More specifically, the subset of epitopes had an average length of 15.45, while the subset of non-epitopes had an average length of 13.97. Among all the epitopes, the peptides with lengths of 16, 15, and 12 amino acids accounted for the largest proportion, i.e., 24.99%, 23.72% and 17.72%, respectively (Figure 1E, 1F). We then analyzed the taxonomic origin of the protein sequences, as provided by the filtered dataset and visualized the distribution of species (Figure 1G, 1H). On the protein level, the curated dataset contained 176 different families. The 21 families with the largest number of epitopes were shown in Figure 1G. The number of epitopes in Bacteria accounted for 16.65%, the Eukaryota for 65.59% and the Viruses for 17.76% of all the proteins, respectively, in the curated dataset. On the epitope-containing region level, the proportions showed differently from the protein level. For example, proportion of viruses was 59.41%, higher than that of the Bacteria (9.17%) and the Eukaryota (31.42%). Overall, the curated dataset had a strong degree of taxonomic diversity.

### Performance of ML methods on benchmarking dataset with sequence and structural features

So far, numerous tools have been developed for xxx. In those tools, a series of sequence or structural features have been adopted. We explored six different conventional ML algorithms, including AB, DT, GB, KNN, LR and RF, using 14 different encoding schemes. For each feature, each algorithm was implemented and optimized using five-fold CV on the training dataset. We repeated five-fold CV ten times by randomly portioning the training dataset. The performances of these 84 ML methods in terms of AUC were shown in **Figure 3**A. The average AUC values of five-fold CV of six ML algorithms ranged from 0.695 (DT) to 0.777 (RF). RF, AB, and LR performed better than other ML-based methods (SGD, KNN, and DT), Next, we studied the average performance for each feature among the six ML methods. The AUC values of five-fold CV ranged from 0.666 (LBA) to 0.768 (AAindexClusterP). Thus, different types of features displayed various accuracies for B cell epitope prediction. We also found that sequence features had superior performance compared to structural features. Due to the limitation of protein structure information, three types of structural features were calculated through computational prediction from protein sequences in this study, and thus, the predicted features might lead to a lower prediction accuracy.

**Figure 3.**
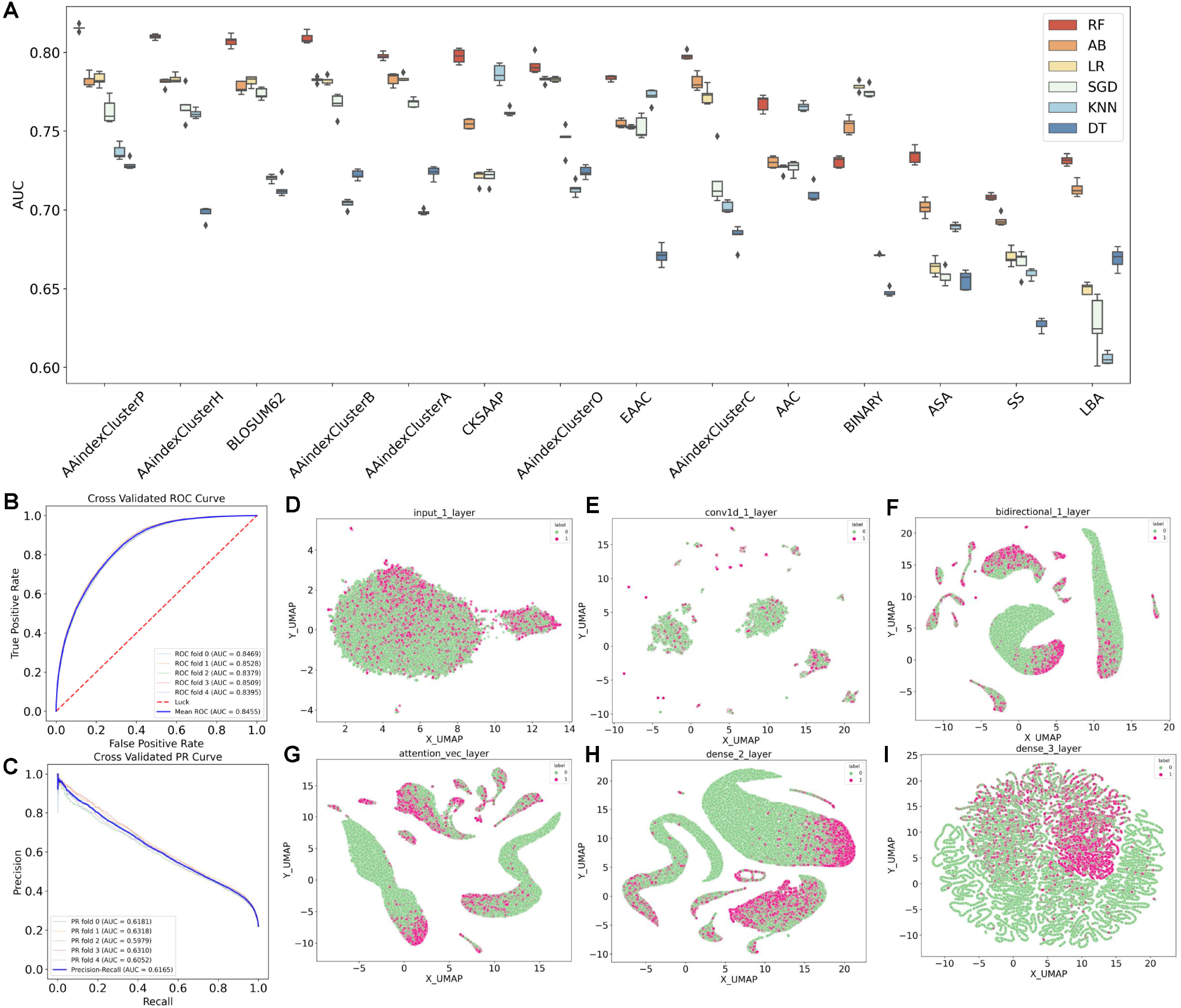
Performance of NetBCE and other machine learning methods. **A.** Performances of 84 machine learning models for the 14 types of features. the area under the receiver operating characteristic (ROC) curve (AUC) values were calculated by five-fold cross-validation. **B-C**. ROC curves (**B**) and (**C**) Precision-Recall (PR) curves for NetBCE by different fold cross-validation. **D-I.** We visualized the epitopes and non-epitopes using the method based on the feature representation at various network layers. Feature representation of different networks indicated discriminative through the network layer hierarchy.

### NetBCE for accurate prediction of linear B cell epitope in proteins

Deep learning has been recently demonstrated to have powerful capability for mining large but complex biomedical data, including image and sequence information extraction and natural language processing. With sufficient B cell assays, the deep neural network can automatically learn informative classification features, making it very appropriate for linear B cell epitope prediction. In this study, a deep learning-based predictor was introduced, called NetBCE, for B cell epitope prediction in the proteins. The NetBCE was implemented with five components: the input layer, convolution-pooling modules, BLSTM layer, attention layer and the output layer. To evaluate the prediction performance of NetBCE, the five-fold CV was performed on the training dataset. The ROC curves were drawn and the corresponding AUC values were calculated. We found that NetBCE had high performance with the average AUC values of 0.846 by five-fold CV, with a range from 0.838 to 0.853 (Figure 3B). Since the numbers of epitopes and non-epitopes were not balanced in the training dataset, we also performed Precision-Recall (PR) analysis and calculated the corresponding AUC values. The PR curve indicates the trade-off between the amount of false positive predictions compared to the amount of false negative predictions. NetBCE achieved average PR AUC values of 0.617 (Figure 3C), suggesting that our model had great potential in predicting functional epitopes with the high precision.

As above, we drew a conclusion that NetBCE was both faithful and robust for the prediction of linear B cell epitope, which might be partly attributed to its deep neural network architecture. NetBCE utilized several excellent deep learning modules, e.g., CNN, LSTM, and Attention, to learn a more efficient and interpretive representation of the epitope sequence hierarchically. To elucidate the capability of hierarchical representation by NetBCE, we visualized the epitopes and non-epitopes using UMAP method based on the feature representation at varied network layers. We found that the feature representation displayed more discriminative along the network layer hierarchy (Figure 3D-I). More specifically, the feature representations for epitopes and non-epitopes sites were mixed at the input layer. As the model continued to train, epitopes and non-epitopes tend to occur in very distinct regions with efficient feature representation.

### Performance evaluation and comparison

To demonstrated the superiority of NetBCE, we first compared the performance of NetBCE with other six ML-based methods (AB, DT, GB, KNN, LR and RF) by AUC value measure. We observed that the AUC values of NetBCE were in a range of 8.77-21.58%, which were generally higher than those of the other six ML-based methods. We further compared NetBCE to three previously developed and available linear B cell epitope predictors, including iLBE, iBCE-EL and BepiPred. Since these three tools did not offer the function for customizing prediction models on other B cell assays, the curated independent dataset was straightly entered to each service to calculate the performance and compare with the prediction result by NetBCE. NetBCE had high performance with the AUC values of 0.840 on the independent dataset (**Figure 4**A). For BepiPred [7], LBtope [9], and iBCE-EL [8] that provide prediction scores for all input, we drew the ROC curves and corresponding AUC values were calculated as 0.688, 0.657 and 0.504, respectively. When compared with the second-best tool BepiPred [7], NetBCE reached an 22.06% AUC value improvement (Figure 4A).

**Figure 4.**
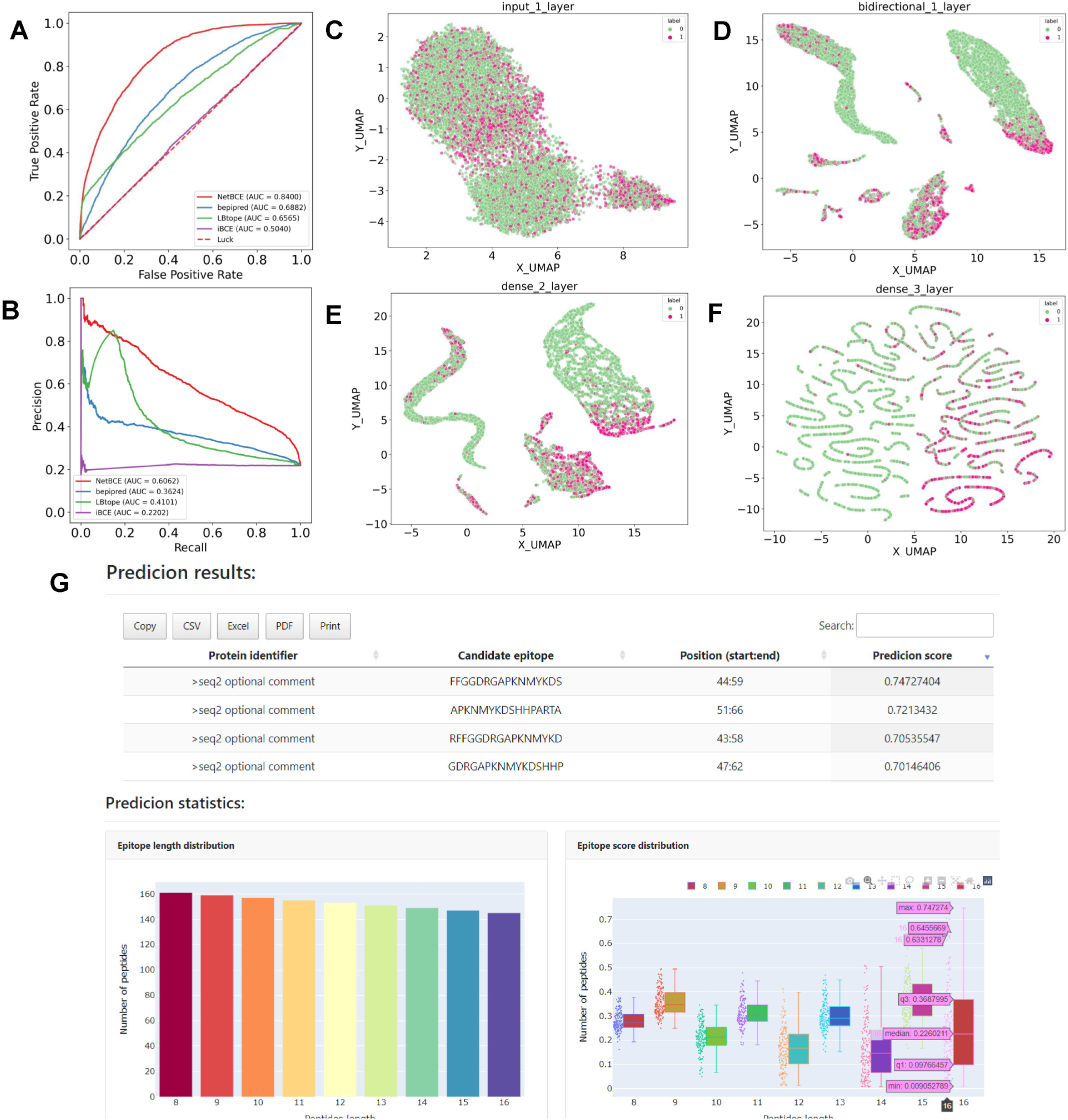
Performance comparison between NetBCE and other tools and NetBCE software. Comparison of NetBCE with other predictors, including BepiPred, LBtope, and iBCE-EL on the independent dataset regarding the (**A**) ROC curves and (**B**) PR curves as well as the corresponding AUC values. **C-F.** The interpretability of the NetBCE on the independent dataset. **G.** NetBCE software. NetBCE provides and visualizes the prediction results in an interactive html file using the Python, PHP, JavaScript and Bootstrap package with an easily readable and interpretable manner.

Moreover, NetBCE reached AUC value (PR) of 0.606 on the independent dataset (Figure 4B), which was superior to other existing tools. To elucidate the underlying mechanism of NetBCE leading to superior performance in the independent dataset, we applied NetBCE to predict the output of important network modules in the model and used UMAP to visualize the predicted feature representation at varied network layers (Figure 4C-F). We found that predicted features became more and more distinguishable with the training of the model. Epitopes and non-epitopes in the independent dataset were mixed at the input layer, culminating with a clear separation in the output layer. In comparison, NetBCE implemented by the interpretable deep learning architecture significantly outperformed other existing tools.

### Usage of NetBCE

Based on to the model constructed in this study, we developed a tool to provide function for linear B-cell epitope prediction. NetBCE provides and visualizes the prediction results in an interactive html file using the Python, PHP, JavaScript and Bootstrap package with an easily readable and interpretable manner. Users can input the candidate proteins in a FASTA format. In addition, user needs to select one or more peptide lengths so that NetBCE can construct a library of candidate epitope peptides. For an example output page in Figure 4G, NetBCE provides a probability score for each candidate peptide with its value in a range from 0 to 1. All prediction results can be copied, printed and downloaded in three formats: “CVS”, “Excel” and “PDF”. NetBCE also provides another two interactive html plots to show the distribution of lengths and scores for all candidate peptides.

## Discussion

In this study, we first compiled an experimentally well-characterized dataset, containing more than 126,000 experimentally linear epitope-containing regions from 3567 protein clusters, through a widely used immunization database (IEDB). Based on the curated benchmark dataset, 14 features were encoded including five sequence-based features, six physicochemical properties-based features and three structural features. All features were evaluated by six conventional machine learning algorithms and the AUC values were calculated through five-fold CV. Our result revealed that predictive power for linear B cell epitope prediction varied greatly by different types of features, but the sequence and physicochemical features had superior performance when compared to structural features. Building on this large data collection, a ten-layer deep learning framework, named NetBCE, was implemented. NetBCE was built by five components: the input layer, convolution-pooling modules, LSTM layer, attention layer and the output layer. To assess the performance of NetBCE, we performed the five-fold CV on the training dataset. NetBCE had high performance with the average AUC values of 0.846, with a range from 0.838 to 0.853, by automatically learn informative classification features. In comparison, NetBCE outperformed conventional ML methods by increasing the AUC value by a range of 8.77-21.58% in same training dataset. Moreover, NetBCE had high performance with the AUC values of 0.840 on the independent dataset, and achieved over 22.06% improvement of AUC value for the linear B cell epitope prediction compared to other tools. Compared to the black box of training process in traditional ML, the interpretability of our model is also easier to explore. To elucidate the capability of hierarchical representation by NetBCE, we visualized the epitopes and non-epitopes based on the predicted feature representation at varied network layers. We found the feature representation came to be more discriminative along the network layer hierarchy, demonstrating our model has excellent classification ability.

In the future, we will continuously strengthen NetBCE by collecting more experimentally identified BCEs into the training dataset. Although the dataset included in the current database is getting larger, a considerable number of BCEs might be false positives that do not have sufficient positive test result. The development of methods for data quality control remains to be a great challenge to minimize the false positives generated by different types of experimental assays. Moreover, more useful features and advanced deep neural network frameworks will be adopted for the development of model for linear B cell epitopes. Taken together, this study reported a novel and accurate approach for the prediction of linear B cell epitopes. We anticipate that the interpretable deep neural network can be easily extended to other sequence-derived prediction task to corroborate much better prediction.

## Code availability

### CRediT author statement

**Haodong Xu:** Conceptualization, Methodology, Software, Data Curation, Visualization, Writing - original draft. **Zhongming Zhao:** Conceptualization, Methodology, Project administration, Supervision, Writing - review & editing, Funding acquisition. All authors read and approved the final manuscript.

## Acknowledgments

The authors thank the members in the Bioinformatics and Systems Medicine Laboratory (BSML) for valuable discussion, and those investigators who generated and shared the reference data. This study was partially supported by National Institutes of Health grants (R01LM012806, R01DE030122, and R01DE029818). We thank the resource support from Cancer Prevention and Research Institute of Texas (CPRIT RP180734 and RP210045). Funding for open access charge: CPRIT (RP180734).

## References

[1] Onda M, Beers R, Xiang L, Lee B, Weldon JE, Kreitman RJ, et al. Recombinant immunotoxin against B-cell malignancies with no immunogenicity in mice by removal of B-cell epitopes. Proc Natl Acad Sci 2011;108:5742–7.

[2] Burger JA, Wiestner A. Targeting B cell receptor signalling in cancer: preclinical and clinical advances. Nat Rev Cancer 2018;18:148–67.

[3] L Dudek N, Perlmutter P, Aguilar I, P Croft N, W Purcell A. Epitope discovery and their use in peptide based vaccines. Curr Pharm Des 2010;16:3149–57.

[4] Potocnakova L, Bhide M, Pulzova LB. An introduction to B-cell epitope mapping and in silico epitope prediction. J Immunol Res 2016;6760830.

[5] Haste Andersen P, Nielsen M, Lund O. Prediction of residues in discontinuous B-cell epitopes using protein 3D structures. Protein Sci 2006;15:2558–67.

[6] Sun P, Guo S, Sun J, Tan L, Lu C, Ma Z. Advances in in-silico B-cell epitope prediction. Curr Top Med Chem 2019;19:105–15.

[7] Jespersen MC, Peters B, Nielsen M, Marcatili P. BepiPred-2.0: improving sequence-based B-cell epitope prediction using conformational epitopes. Nucleic Acids Res 2017;45:W24–W9.

[8] Manavalan B, Govindaraj RG, Shin TH, Kim MO, Lee G. iBCE-EL: a new ensemble learning framework for improved linear B-cell epitope prediction. Front Immunol 2018;9:1695.

[9] Singh H, Ansari HR, Raghava GP. Improved method for linear B-cell epitope prediction using antigen’s primary sequence. PloS One 2013;8:e62216.

[10] Zobayer N, Hossain AA, Rahman MA. A combined view of B-cell epitope features in antigens. Bioinformation 2019;15:530.

[11] El-Manzalawy Y, Dobbs D, Honavar V. Predicting flexible length linear B-cell epitopes. Comput Syst Bioinformatics Conf, 2008; 7:121–32.

[12] Emini EA, Hughes JV, Perlow D, Boger J. Induction of hepatitis A virus-neutralizing antibody by a virus-specific synthetic peptide. J Virol 1985;55:836–9.

[13] Ning W, Xu H, Jiang P, Cheng H, Deng W, Guo Y, et al. HybridSucc: a hybrid-learning architecture for general and species-specific succinylation site prediction. Genom Proteom Bioinform 2020;18:194–207.

[14] Xu H-D, Liang R-P, Wang Y-G, Qiu J-D. mUSP: a high-accuracy map of the in situ crosstalk of ubiquitylation and SUMOylation proteome predicted via the feature enhancement approach. Brief Bioinform 2021;22:bbaa050.

[15] Xu H, Jia P, Zhao Z. DeepVISP: deep learning for virus site integration prediction and motif discovery. Adv Sci 2021;8:2004958.

[16] Xu H, Jia P, Zhao Z. Deep4mC: systematic assessment and computational prediction for DNA N4-methylcytosine sites by deep learning. Brief Bioinform 2021;22:bbaa099.

[17] Senior AW, Evans R, Jumper J, Kirkpatrick J, Sifre L, Green T, et al. Improved protein structure prediction using potentials from deep learning. Nature 2020;577:706–10.

[18] Wang C, Xu H, Lin S, Deng W, Zhou J, Zhang Y, et al. GPS 5.0: an update on the prediction of kinase-specific phosphorylation sites in proteins. Genom Proteom Bioinform 2020;18:72–80.

[19] Vita R, Mahajan S, Overton JA, Dhanda SK, Martini S, Cantrell JR, et al. The immune epitope database (IEDB): 2018 update. Nucleic Acids Res 2019;47:D339–D43.

[20] Ning W, Jiang P, Guo Y, Wang C, Tan X, Zhang W, et al. GPS-Palm: a deep learning-based graphic presentation system for the prediction of S-palmitoylation sites in proteins. Brief Bioinform 2021;22:1836–47.

[21] Kawashima S, Pokarowski P, Pokarowska M, Kolinski A, Katayama T, Kanehisa M. AAindex: amino acid index database, progress report 2008. Nucleic Acids Res 2007;36:D202–D5.

[22] Sun P, Yu Y, Wang R, Cheng M, Zhou Z, Sun H. B-cell Epitope prediction method based on deep ensemble architecture and sequences. 2019 IEEE International Conference on Bioinformatics and Biomedicine (BIBM) 2019:94–7.

[23] Yang Y, Heffernan R, Paliwal K, Lyons J, Dehzangi A, Sharma A, et al. SPIDER2: a package to predict secondary structure, accessible surface area, and main-chain torsional angles by deep neural networks. Prediction of protein secondary structure. Springer, 2017, 55–63.

[24] Min S, Lee B, Yoon S. Deep learning in bioinformatics. Brief Bioinform 2017;18:851–69.

[25] McInnes L, Healy J, Melville J. Umap: Uniform manifold approximation and projection for dimension reduction. arXiv preprint arXiv:1802.03426 2018.

[26] Geer LY, Marchler-Bauer A, Geer RC, Han L, He J, He S, et al. The NCBI biosystems database. Nucleic Acids Res 2010;38:D492–D6.

[27] Consortium U. UniProt: a worldwide hub of protein knowledge. Nucleic Acids Res 2019;47:D506–D15.

[28] Fu L, Niu B, Zhu Z, Wu S, Li W. CD-HIT: accelerated for clustering the next-generation sequencing data. Bioinformatics 2012;28:3150–2.

[29] Pang Y, Sun M, Jiang X, Li X. Convolution in convolution for network in network. IEEE Trans Neural Netw Learn Syst 2017;29:1587–97.

[30] Huang Z, Xu W, Yu K. Bidirectional LSTM-CRF models for sequence tagging. arXiv preprint arXiv:1508.01991 2015.

[31] Wang F, Jiang M, Qian C, Yang S, Li C, Zhang H, et al. Residual attention network for image classification. Proc IEEE Comput Soc Conf Comput Vis Pattern Recognit 2017:3156–64.

[32] Bergstra J, Komer B, Eliasmith C, Yamins D, Cox DD. Hyperopt: a python library for model selection and hyperparameter optimization. Comput Sci Discov 2015;8:014008.

